# Rapid Development of Neutralizing and Diagnostic SARS-COV-2 Mouse Monoclonal Antibodies

**DOI:** 10.1101/2020.10.13.338095

**Authors:** Asheley P. Chapman, Xiaoling Tang, Joo R. Lee, Asiya Chida, Kristina Mercer, Rebekah E. Wharton, Markus Kainulainen, Jennifer L. Harcourt, Roosecelis B. Martines, Michelle Schroeder, Liangjun Zhao, Anton Bryksin, Bin Zhou, Eric Bergeron, Brigid C. Bollweg, Azaibi Tamin, Natalie Thornburg, David E. Wentworth, David Petway, Dennis Bagarozzi, M.G. Finn, Jason M. Goldstein

**Affiliations:** School of Chemistry and Biochemistry, Georgia Institute of Technology, 901 Atlantic Dr., Atlanta, GA, USA, 30306; Immunodiagnostic Development Team/ Reagent Diagnostic Services Branch (RDSB)/DSR/NCEZID/CDC, 1600 Clifton Rd NE. Atlanta, GA, USA, 30333; Vaccine Preparedness Team/ Virology Surveillance and Diagnosis Branch (VSPB)/ID/NCIRD/CDC, 1600 Clifton Rd NE. Atlanta, GA, USA, 30333; Viral Special Pathogens Branch VSPB/DHCPP/NCEZID/CDC, 1600 Clifton Rd NE. Atlanta, GA, USA, 30333; Respiratory Disease Branch (RDB)/DVD/NCIRD/CDC, 1600 Clifton Rd NE. Atlanta, GA, USA, 30333; Infectious Disease Pathology Branch (IDPB)/DHCPP/NCEZID/CDC, 1600 Clifton Rd NE. Atlanta, GA, USA, 30333; Parker H. Petit Institute for Bioengineering and Bioscience, Georgia Institute of Technology, Atlanta, GA, USA, 30306; Division of Laboratory Sciences/NCEH/CDC, 4770 Buford Hwy, Atlanta, GA 30341, USA; School of Biological Sciences, Georgia Institute of Technology, 901 Atlantic Dr., Atlanta, GA, USA, 30306

**Author notes:** Authors provided equivalent contributions (Xiaoling Tang, Joo R. Lee, Asiya Chida).

## Abstract

The need for high-affinity, SARS-CoV-2-specific monoclonal antibodies (mAbs) is critical in the face of the global COVID-19 pandemic, as such reagents can have important diagnostic, research, and therapeutic applications. Of greatest interest is the ~300 amino acid receptor binding domain (RBD) within the S1 subunit of the spike protein because of its key interaction with the human angiotensin converting enzyme 2 (hACE2) receptor present on many cell types, especially lung epithelial cells. We report here the development and functional characterization of 29 nanomolar-affinity mouse SARS-CoV-2 mAbs created by an accelerated immunization and hybridoma screening process. Differing functions, including binding of diverse protein epitopes, viral neutralization, impact on RBD-hACE2 binding, and immunohistochemical staining of infected lung tissue, were correlated with variable gene usage and sequence.

---

Previous beta-coronavirus outbreaks from SARS-CoV (2003) and MERS-CoV (2008) highlighted the value of novel antibody development in the rapid diagnosis of infection. Studies of these earlier coronaviruses have informed current SARS-CoV-2 vaccine design, especially concerning epitope motifs to target for virus neutralization. The ~300 amino acid receptor binding domain (RBD) within the S1 subunit of the spike protein (Fig. 1a) is particularly significant, as it contains the five contact residues (L455, F486, Q493, S494, N501)^1,2^ previously shown to be important for viral entry through interactions with the ACE2 receptor present on many cell types, especially lung epithelial cells.^3^

**Figure 1.**
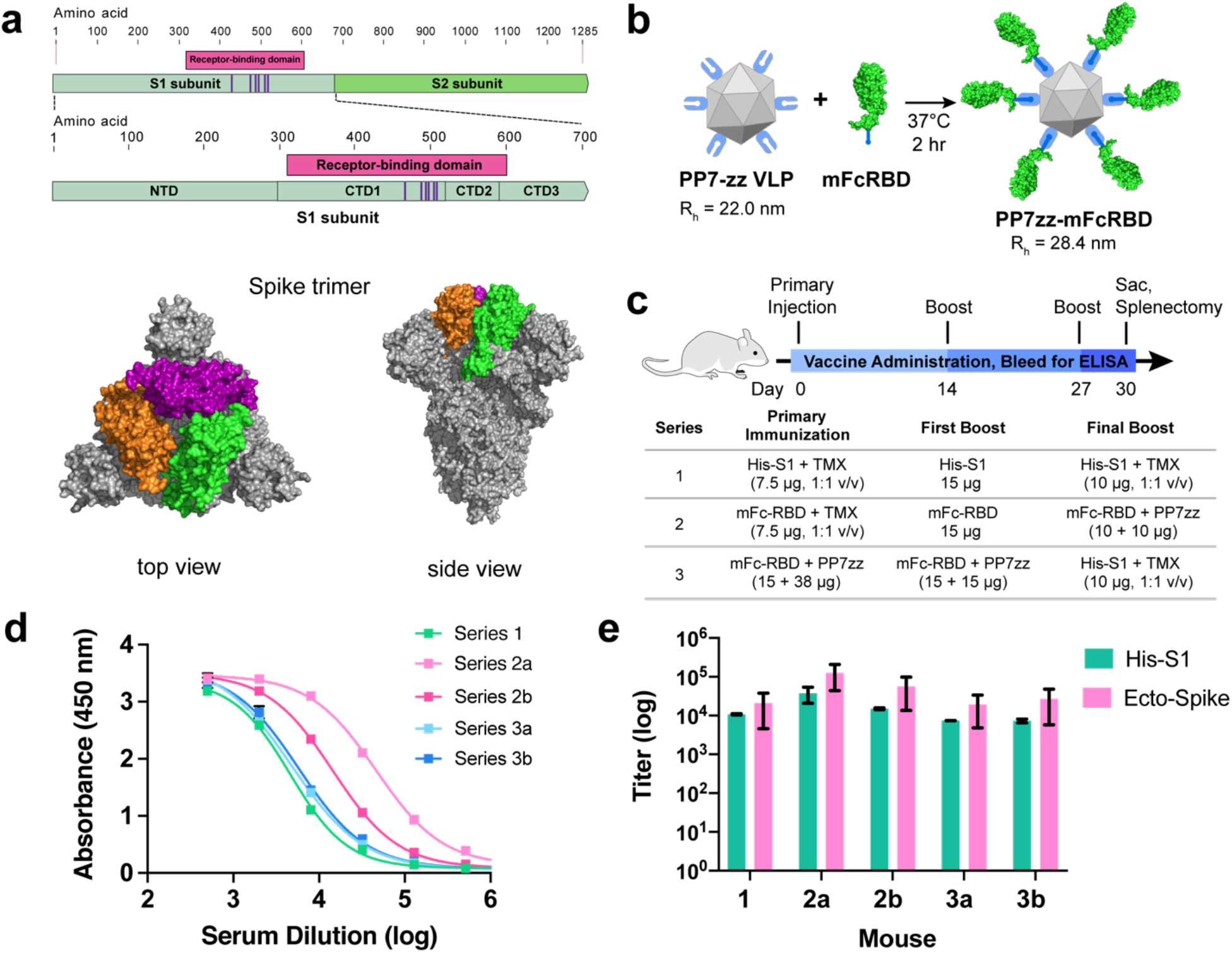
SARS-CoV-2 Spike protein subunit vaccine strategy and humoral immune response in mice. **a**, Recombinant spike subunit 1 protein (His-S1, residues 1-681) or S1 Receptor Binding Domain (mouse Fc-RBD, residues 319-541, ACE2 contact residues in purple; PDBID: 6vxx) antigens. **b**, VLP display of Fc-tagged antigens using the PP7 particle bearing 120 ZZ-domains; a 1:1 mass ratio of mFc-RBD and VLP provides a Fc:ZZ molar ratio of approximately 0.8. Rh = hydrodynamic radius measured by dynamic light scattering in phosphate buffer. **c**, Vaccine schedule and strategy. Six-week old female BALB/c mice (n = 3 per group) were immunized with primary antigen and adjuvant on day 0 followed by boosts on days 14 and 27. Blood was collected for ELISA on days 0, 14, 21 and 30. **d**, ELISA responses for serum dilutions against plated His-S1 protein from the sacrificed mice at day 30. **e**, Titer values from ELISA analysis as in panel d, against plated His-S1 or spike ectodomain protein. Immunization series defined in panel c; a and b designate different mice within that series. Experimental error represents standard deviation.

In order to target this key polypeptide motif, we employed three different immunogens: a commercially-available S1 domain (residues 1-681)^4^ with C-terminal histidine tag (designated His-S1), a commercial RBD sequence (residues 319-541^4^) fused at the C-terminus with a mouse IgG1 Fc domain (designated mFc-RBD) (Fig 1a), and the noncovalent complex of mFc-RBD with the PP7 bacteriophage virus-like particle (VLP) engineered to express two sequential Fc-binding Z-domains at 120 places on the VLP exterior surface (designated PP7zz).^5^ The last construct provided a large (Fig. 1b) polyvalent display of receptor binding domains which we hoped would productively engage immune cells and stimulate affinity maturation. The use of Fc-tagged antigens is well known to increase antigen uptake and processing,^6^ including many examples with viral antigens.^7–11^ While the virus-like particles used here were regarded as self-adjuvanting,^12,13^ the recombinant S1 and RBD proteins were usually augmented by TiterMax® Gold emulsion adjuvant. Three sets of BALB/c mice were immunized with different accelerated vaccination schedules (Fig. 1c), characterized by the use of only the recombinant His-S1 domain (Series 1), a combination of mFc-RBD and PP7zz-displayed mFc-RBD immunogens beginning with mFc-RBD (Series 2), and the VLP-displayed mFc-RBD with a final focusing boost of His-S1 (Series 3). Individual mice or pairs of mice were chosen for maximum immune response as indicated by serum antibody titer (Fig. 1d,e), rather than larger cohorts as would be necessary to determine reproducibility and mechanistic trends.

Splenocyte harvest (day 30) and myeloma fusion (Extended Data Fig. 2a, b) were followed by robotic selection of high IgG secretors from 3D semi-solid culture. Enzyme-linked immunosorbent assay (ELISA) analysis of supernatants from individual cultures of these cells against immobilized antigenic protein (recombinant His-S1 or engineered SARS-CoV-2 spike ectodomain protein,^14,15^ presumed to exist as monomer and noncovalent trimer, respectively) revealed the series 2 immunization protocol featuring mFc-RBD primary immunization to be superior in producing SARS-CoV-2 relevant antibodies (Fig. 2a, Extended Data Fig. 2b). Each of 159 spike-binding mAbs selected by ELISA (157 from series 2, two from series 3) were further characterized by semi-quantitative measurement of the binding strength of purified antibody or hybridoma supernatant to soluble spike ectodomain using biolayer interferometry (BLI). Twelve of these showed apparent avidity (reported as inverse of the adsorption constant, 1/K_ads_) better than 1 nM and seventeen exhibited 1/K_ads_ values between 1 and 10 nM (Fig. 2b, Table 1). Four mAbs demonstrating markedly lower avidity by BLI (3D2, 3G6, 3B1, 3A6) were also carried forward based on their demonstration of interesting or unusual properties as described below. All of the 33 selected mAbs were IgG1 (k light chain), showed strong binding to both plated S1 subunit and spike ectodomain proteins, with overall larger signals for the latter under otherwise identical ELISA conditions (Figure 2c, Table S1), and bound (in hybridoma supernatant) inactivated (gamma-irradiated) SARS-CoV-2 virus (Fig. 2d). The binding of serum, supernatants, and antibodies to the ectodomain were <u>not</u> found to be consistently affected by storage or incubation of the target protein at various temperatures, a concern raised by a recent report that the structure of the ectodomain may be cold-sensitive (Extended Data Fig. 3).^16^

**Figure 2.**
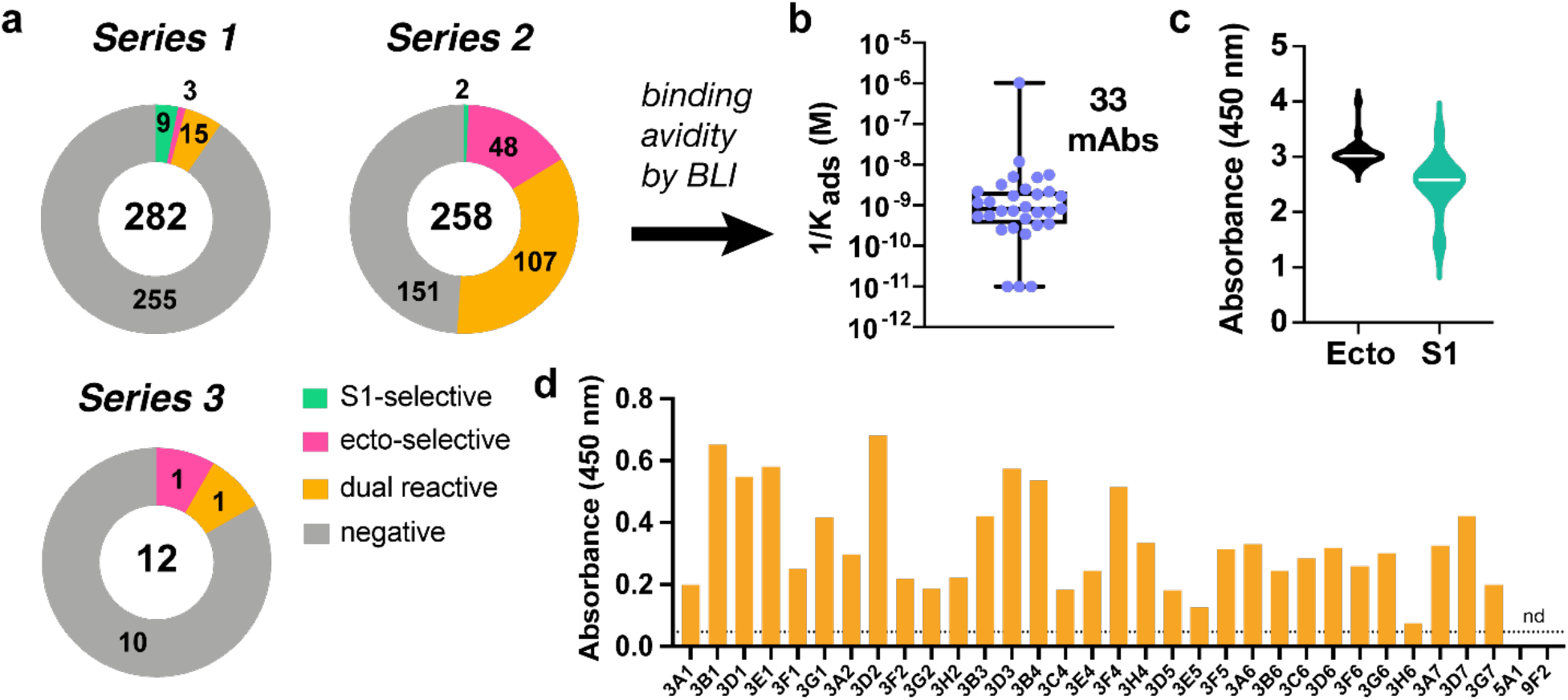
Characterization of anti-SARS-COV-2 antibodies as supernatants from hybridoma clones. **a**, Antigen specificity of antibody binding from IgG-secreting hybridomas chosen from Series 1-3 immunization schedules. Numbers inside circles are total clones selected, numbers within each portion reflect binding selectivity. **b**, Distribution of affinities of the 33 selected clones as measured by BLI. **c**, Binding to two different Spike recombinant proteins (ecto Spike trimer and S1 polypeptide) as determined by ELISA. **d**, Binding of antibodies to γ-irradiated WA-SARS-CoV-2 by ELISA (15 ng/mL virus plated, supernatant undiluted, secondary antibody = goat anti-mouse IgG HRP conjugate).

**Table 1.**
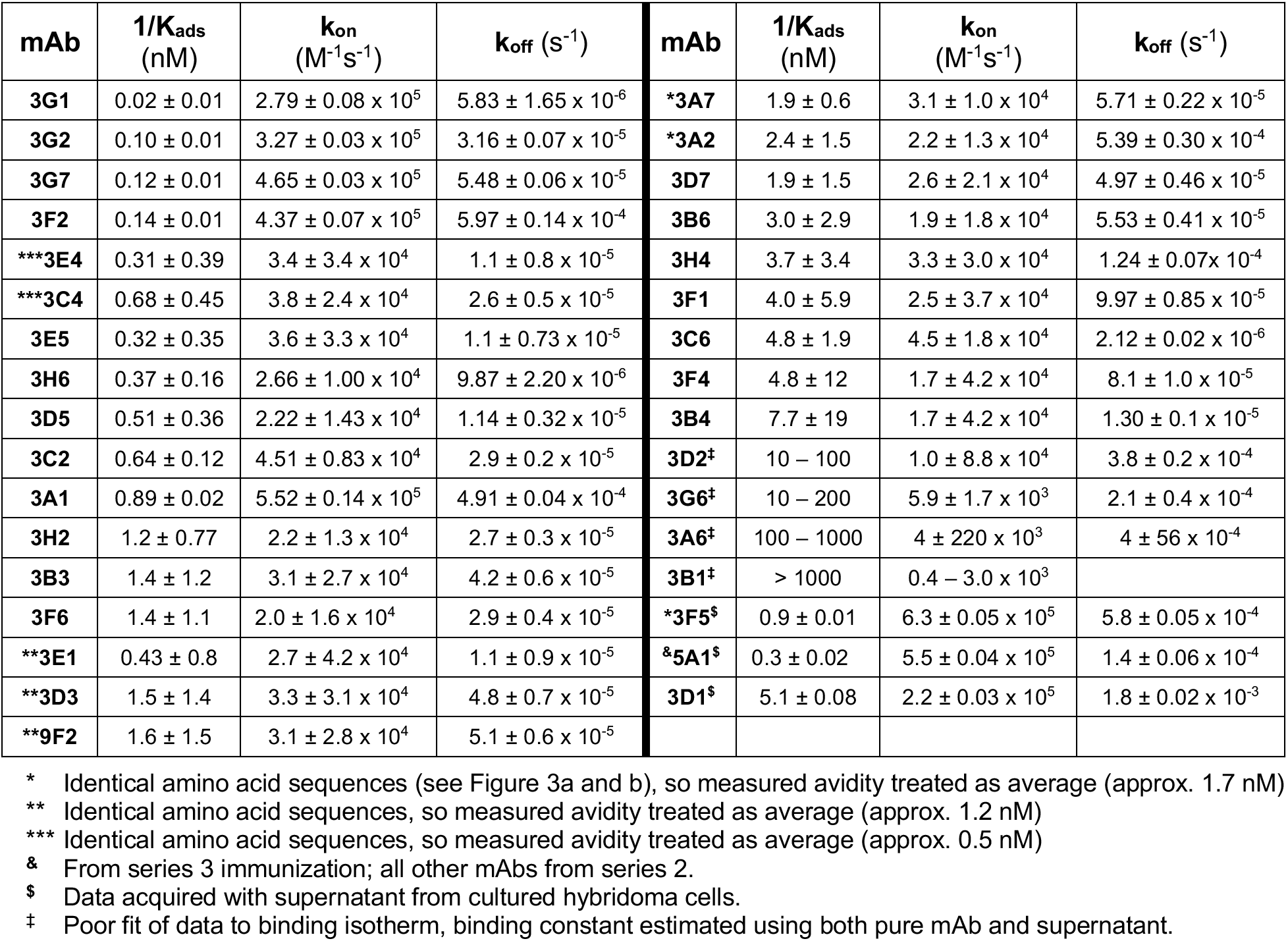
Binding parameters of selected antibodies. Kinetic and thermodynamic parameters for binding of soluble spike ectodomain protein to antibodies immobilized by capture on anti-mouse Fc-coated biolayer interferometry sensor chips (ForteBio Octet).

Variable-region sequencing showed three sets of identical mAbs: 3A2/3A7/3F5 (plus 3C6, identical in the heavy chain but with some variations in the light chain, corresponding to modest functional differences as described below); 3C4/3E4, and 3E1/3D3/9F2. Since the spatial separation of cells in the 3D culture matrix usually far exceeded the resolution of the clone-picking apparatus, it is unlikely that these sets of identical antibodies arose from the same hybridomas. (The lone exception was 3E1 and 9F2, which originated from same clone sampled twice, Extended Data Fig 2b.) Overall, the sequences of these high-affinity antibodies clustered into several discreet groups based on heavy- or light-chain similarities (Fig. 3a, b). The most diverse region in terms of length and amino acid composition proved to be heavy-chain CDR3 (Fig. 3c, d), whereas greater diversity in gene usage was observed in the light chain (Fig. 3e). Gene clustering was found to have relevance to functions described below. Thus, neutralization (mAb groups 1-3) was associated primarily with the presence of IGHV1 and 8, and with IGKV1, 3, and 6; the large non-neutralizing Group 3 all shared the IGHV14-1 gene and about half also incorporated IGKV6-32.

**Figure 3.**
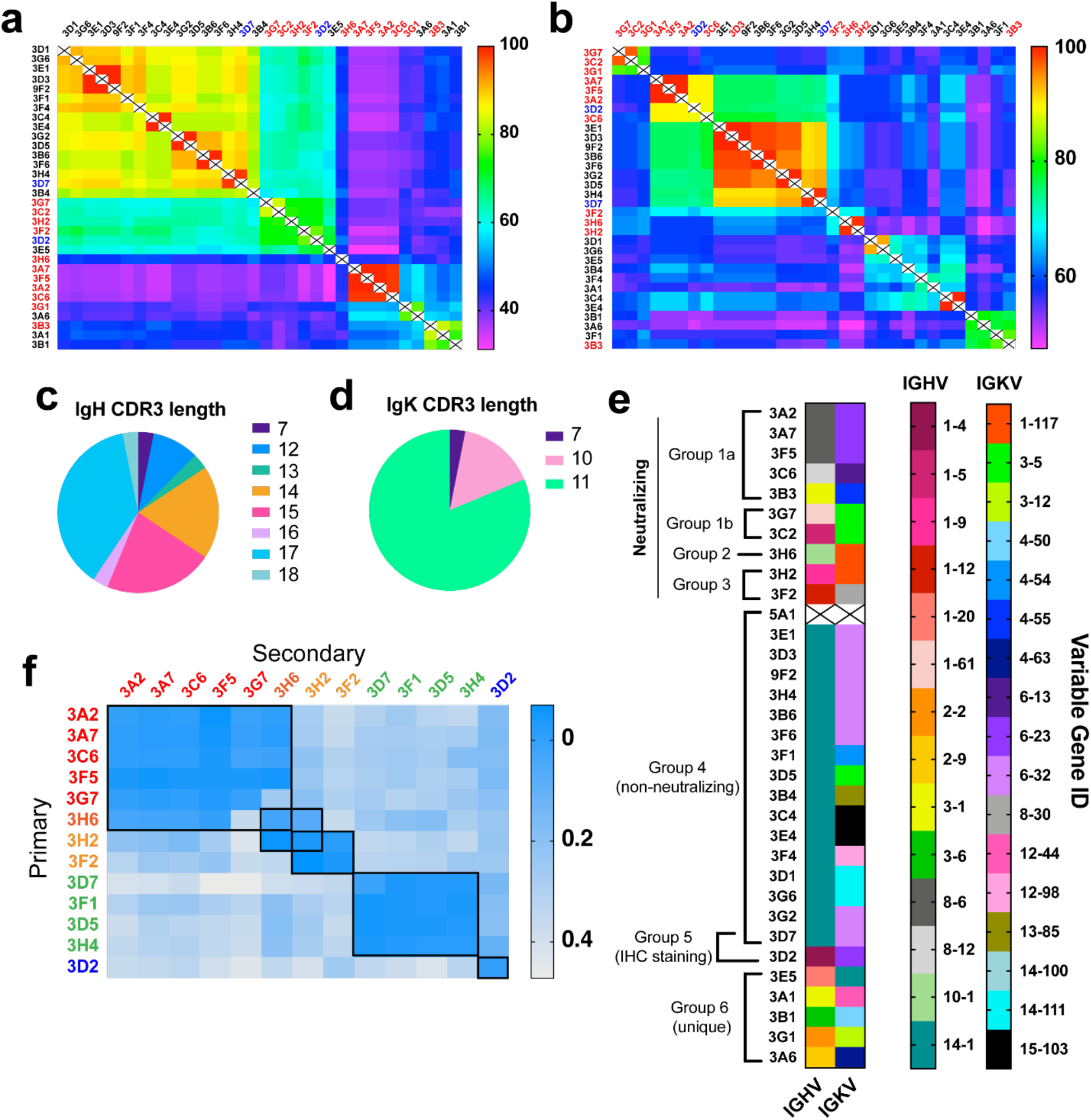
SARS-CoV-2 mAb sequences and epitope binning. **a,b**, Overall similarity of (**a**) heavy chain and (**b**) light chain anti-SARS-CoV-2 antibody sequences, scored by percent of the total number of identical amino acids at each position. Antibody sequences were analyzed via MiXCR software and sequence alignment performed by Geneious. Neutralizing (red) and IHC staining mAbs (blue) determined via functional assays (Fig. 4) **c,d**, Distribution of CDR3 amino acid length in heavy (**c**) and light (**d**) chains of 32 SARS-COV-2 mAbs. **e**, IGHV and IGKV genes identified for mAbs from sequences in panels (**a**) and (**b**). mAb 5A1 was not sequenced. Antibody grouping includes sequence, epitope, and function-level characteristics. **f**, Representative assessment of competitive mAb binding to recombinant ectodomain immobilized to streptavidin-coated BLI sensor chips via *Strep-tag®* sequence. Binding to the “primary” antibody was validated by BLI, followed by washing and exposure to the secondary antibody to generate the binding signal of interest.

The diversity of epitopes recognized in binding the spike ectodomain was assessed by pairwise competitive binding measurements using BLI (Fig. 3f). Low values indicated competition of the two antibodies for identical, nearby, or allosterically-related epitopes, whereas large BLI signals indicated non-competitive antibody association to different epitopes. Six major binding groups were thereby identified (Fig. 3e, Extended Data Fig. 5-7). Some correlation was observed with sequence overlap (as shown by the group labels in Fig. 3a, b), but a much greater degree of correlation was observed to heavy-chain sequences and specific IGHV and IGKV combinations (noted above and Extended Data Fig. 5), and to other functional assays.

The effect of the selected mAbs on the binding of the SARS-CoV-2 RBD to the human ACE2 receptor was determined by two *in vitro* methods: a direct binding measurement using both longer (residues 1-740 = nearly the full polypeptide, referred to here as the “full-length” protein) and shorter (residues 1-614, lacking the C-terminal collectrin-like domain, termed “truncated”) ACE2 constructs, and a split luciferase assay employing the truncated version (Fig. 4a, b). Ten mAbs (3A2/3A7/3F5 = identical sequence, 3C6, 3B3, 3H6, 3H2, 3F2, 3G1, and 3A6) were found to strongly inhibit RBD binding to both versions of hACE2 in dose-dependent fashion. Eight of these antibodies (3A2/3A7/3F5, 3B3, 3H6, 3H2, 3F2, and 3G1) plus two others (3G7, 3C2, also inhibitors of the RBD-ACE2 interaction but with a different profile, discussed below) exhibited potent neutralization of SARS-CoV-2-WA P#4, an early patient isolate,^17^ in cell culture (Figure 4c, Extended Data Table S1). These results were supported by a second neutralization assay employing a GFP-expressing variant of SARS-CoV-2^18^ (Fig. 4c).

**Figure 4.**
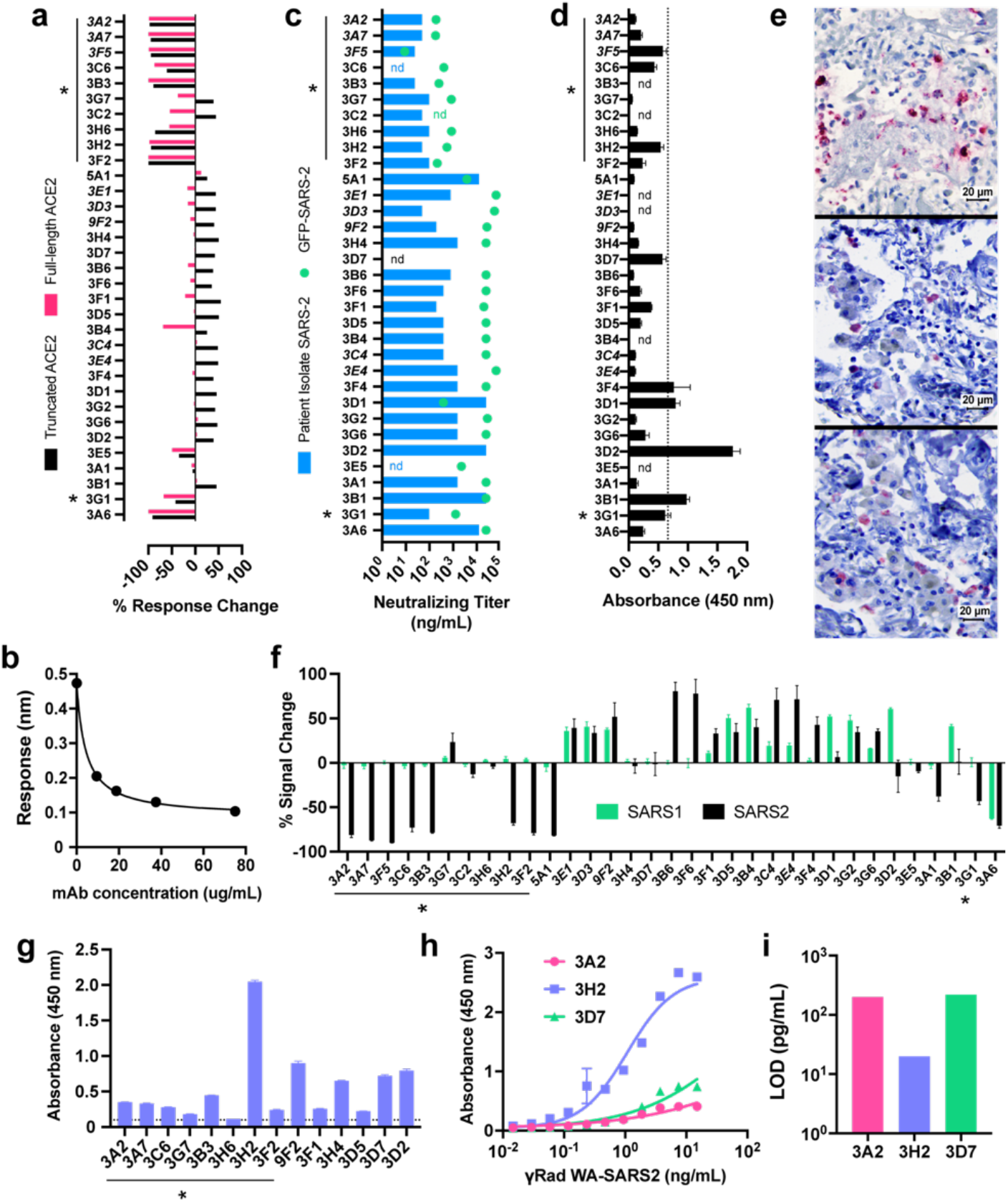
Functional utility of SARS-CoV-2 mAbs. **a**, Assessment of inhibition of S1 RBD binding to human ACE2 receptor as measured by BLI, performed at a single antibody concentration. Both full-length ACE2 (SinoBio, residues #1-740, pink bars) and truncated ACE2 (residues #1-614, black bars) were included. Data represents percent of signal change normalized to response of ACE2/RBD binding with no mAb present, bars represent standard deviation of technical replicates. Neutralizing antibodies in both assays are denoted by asterisk; adjacent mAbs in italics are identical. **b**, Representative (mAb 3G7) dose-dependent antibody-mediated inhibition of ecto-Spike binding to fulllength human ACE2 receptor as measured by BLI; readout is the wavelength shift in nm in the interferometric response.^20,21^ **c**, *In vitro* mAb neutralization of WA-SARS-CoV-2 virus (blue bars) or of GFP-expressing SARS-CoV-2 (green dots). Asterisks mark particularly effective entries. **d**, Binding of antibodies to biotinylated peptide (amino acids 486-501 with C488S substitution; containing four out of five reported RBD-hACE2 contacts) plated on streptavidin-coated plates as measured by ELISA. Dashed line represents the average plus twice the standard deviation of the values obtained for replicates involving three identical mAbs (3A2, 3A7, 3F5); error bars = standard deviation. **e**, Immunohistochemical staining of lung tissue from an early confirmed SARS-CoV-2 fatal case by (*top*) SARS polyclonal nucleocapsid antibody previously validated as an effective reagent,^22^ *(middle)* mAbs 3D2, and *(bottom)* 3D7 (following paraffin removal, rehydration, and heat-induced antigen retrieval, 1:100 mAb dilution, antigen detection appears in red). **f**, SARS specificity of mAbs determined by split nanoluciferase assay including either SARS1 or SARS2 RBD incubated with mAb followed by ACE2. Percent change is normalized to ACE2/RBD signal without antibody. Negative values are interpreted as inhibition, positive as an allosteric binding enhancement observed only with truncated ACE2. **g**, ELISA detection of mAb binding to gamma-inactivated SARS-COV-2 (1 μg/mL TCID_50_ units plated). The dotted line marks the magnitude of background signal. **h**, dose-dependent gamma-inactivated SARS-COV-2 binding of select mAbs (5 μg/mL mAb) measured by ELISA, sigmoidal dose-response curve fitted by GraphPad Prism8. **i**, Limit of detection quantification of mAbs in panel g. LOD calculated as the minimum virus concentration giving rise to a signal equivalent to the mean background signal plus 3 experimental standard deviations (GraphPad Prism 8).

Twenty-one other mAbs exhibited significant levels of apparent RBD/hACE2 antibody-dependent binding enhancement with the truncated ACE2 receptor, but 17 of these had little or no effect on the interaction of RBD with full-length ACE2; the remaining 3 (3G7, 3C2, 3B4) showed moderate levels of full-length ACE2-RBD binding inhibition (Fig. 4a, f). Since the truncated ACE2 lacks the receptor’s dimerization domain,^19^ it is clear that these antibodies recognize different epitopes, and are distinct in functional class, from the strong inhibitors of RBD binding to both hACE2 constructs.

The *in vitro* split luciferase functional assay mirrored these binding results with several interesting exceptions. Thus, 3H6, 3G7, and 3C2 had strong effects on RBD-ACE2 direct binding but little influence on split luciferase readout; contrariwise, 3A1 and 5A1 inhibited the SARS-2 split luciferase reporter but did not inhibit direct binding. As described below, these functional oddities were matched by independently observed differences in competitive epitope binding. When the RBD component of the split luciferase assay was changed to the SARS-CoV-1 sequence, all but one strongly neutralizing antibody was shown to be selective for SARS-CoV-2 (Fig. 4f). [One neutralizer (3H6) had no effect on the split luciferase assay, and one strong ACE2 binding inhibitor (3A6) was not neutralizing. The latter, interestingly, was the only mAb to inhibit RBD binding to both CoV-1 and CoV-2 ACE2 sequences and also had low overall binding affinity for the recombinant spike ectodomain protein.] In contrast, of the 17 antibodies eliciting a strengthened split luciferase signal, 14 gave roughly comparable results for both SARS-CoV-1 and CoV-2, while only three were selective for each variant (3B6, 3F6, and 3F4 for CoV-2; 3D1, 3D2, and 3B1 for CoV-1).

Based on sequence variations and epitope binning experiments (Fig 3), we conclude that the groups [3A2/3A7/3F5, 3C6, 3B3], [3G7, 3C2], [3H6], and [3H2, 3F2] recognize at least four different epitopes of the SARS-CoV-2 receptor binding domain, the general target for neutralizing human antibodies as well.^23^ Four mAbs (3B1, 3D2, 3F4, 3D1) were found to bind to a detectable degree to a synthetic biotinylated 15-mer peptide of the receptor binding motif covering amino acids 486-501 (Fig. 4d), which contains four out of five key RBD-hACE2 contact residues.^1,2^ None of these four antibodies are strongly neutralizing, suggesting that the linear (and presumably unstructured) peptide is not presented by the infectious virus.

Such a peptide motif, however, can apparently be presented when the virion is degraded. Among our high-affinity mAb panel, we identified two (3D2 and 3D7) from a subset of 13 (including 3G7, 3F2, 5A1, 3A1, 3F1, 3B1, 3A6, 3G6) that preferentially stained SARS-CoV-2 viral spike protein in lung tissue from infected patients. These samples underwent standard fixation and subsequent heat treatment to expose antigen, likely in denatured form, at the surface (Fig. 4e). As noted above, these clones have functional characteristics opposite to those desired of a therapeutic molecule: they exhibited little or no detectable neutralization activity and enhanced binding of spike protein to the truncated ACE2 receptor. Heat-inactivated virions obtained from ATCC were recognized by ELISA by only 3H2 (Extended Data Fig. 4c) among the mAbs described here.

Recognition of gamma-irradiated SARS-CoV-2-WA virions was also not uniform among mAbs having roughly equal avidities for the recombinant spike ectodomain protein, as illustrated by ELISA analysis of 15 antibodies in Fig. 4g. Representative dose-response analysis showed that, while both 3H2 and 3A2 were equivalently potent in *in vitro* neutralization, the former mAb generated significantly larger ELISA signals at equivalent concentrations (Fig. 4h). This resulted in a limit of detection of gamma-inactivated particles using 3H2 that was lower by an order of magnitude (Fig. 4i). While it is sometimes assumed that gamma-irradiation does not alter viral surface antigens,^24^ this is not at all certain.^25^ We are aware of no studies of this question for coronavirus.

In summary, immunization using Fc-domain (and virus-like particle assisted) display of the SARS-CoV-2 receptor binding domain, followed by rapid parallel hybridoma production and screening, provided a panel of binding agents with diverse and useful functions in the context of virus binding and infectivity. This approach is complementary to that of teams led by Crowe and Walker, who isolated large panels of human anti-SARS-CoV-2 mAbs from B cells from infected patients.^26,27^ As in those reports, a large number (here, 29 out of 33) anti-SARS-CoV-2 monoclonal antibodies were obtained exhibiting low-nanomolar or better avidity for the target spike ectodomain, consistent with many other reports validating this protein as a viable immunological target. We identified three groups of neutralizing antibodies recognizing distinct epitopes on the RBD motif as inferred from sequence and competitive binding experiments (Fig. 3e, f; Extended Data Fig. 6, 7). A fourth group, the largest in number, were not strongly neutralizing and had either no effect on, or enhanced in one assay, the recognition of human ACE2 by the recombinant RBD. A fifth functional group stained fixed and heat-treated lung tissue from infected COVID-19 patients. Each *functional* class was also largely distinctive in its epitope recognition and in its sequence.

## Methods

### General

Monoclonal antibody sequencing and functional tests were performed by blinded researchers. All data were processed and graphed using GraphPad Prism v.8.

### Recombinant ectodomain protein

We employed the stabilized prefusion spike construct of Graham and McLellan (designated “s2p” by those investigators, and “recombinant spike ecto” (r-spike ecto) here.^15^ This polypeptide lacks the transmembrane sequence of the natural virus, has a mutated furin cleavage site (RRAR to GSAS) to block proteolytic processing, and is capped at the C-terminus with a His8 tag followed by two Strep-Tag® II sequences (WSHPQFEK) bridged by a flexible connector (GGGSGGGGSGGSA).

### Expression and purification of His-tagged recombinant SARS-CoV-2 spike S1 and ectodomain proteins

Expi293F™ cells (Thermo Fisher Scientific) were cultured on an orbital shaker platform (~125 rpm) in Expi-293 expression medium at 37°C with 8% CO2. On the day of transient transfection, 400 μg of pCDNA3.1-SARS-CoV2-S1 recombinant plasmid DNA (provided by Dr. Lisa Mills at the Centers for Disease Control and Prevention) was mixed with ExpiFectamine™ 293 reagent in Opti-MEM™ I reduced serum medium and transfected into 400 mL of Expi293F cells following the manufacturer’s instructions. About 20 h post-transfection, ExpiFectamine™ 293 Transfection Enhancer 1 and Enhancer 2 were added to the flask containing transfected cells. On day 6 posttransfection, cells were harvested and centrifuged at 5,000 x g for 30 minutes at 4°C. The supernatant was filtered through a 0.22-μM filter and loaded onto a 5 mL HisTrap FF crude column (GE) for purification. The column was washed with buffer A (20 mM Tris-HCl, 500 mM NaCl, 40 mM imidazole, pH 7.5). His-tagged SARS-CoV-S1 recombinant protein was then eluted with buffer B (20 mM Tris-HCl, 500 mM NaCl, 500 mM imidazole, pH 7.5) and dialyzed into 0.01M PBS, pH 7.4.^28^ To produce the ectodomain protein, an Expi293F™ stable cell line was constructed to express r-spike ecto. Six liters of these cells were seeded at 5×10^5^ cells per mL into flasks and cultured at 37°C and 8% CO2 with orbital shaking (~125 rpm). After 7 days of incubation, cells were harvested and centrifuged at 5000 x g for 30 minutes at 4°C. The supernatant was filtered through 0.22-μM filter and loaded onto 5 mL HisTrap FF crude column (GE) for purification as described above.

### Immunizations, Antigens, and B cell extraction

Six-week old female pathogen-free BALB/c mice were obtained from Charles River Laboratories (Wilmington, MA) and housed in the Physiological Research Laboratory at the Georgia Institute of Technology. All animal protocols and procedures were approved by the Georgia Tech Institutional Animal Care and Use Committee. Mice were immunized subcutaneously on both sides of lower anterior abdomen on day 0 with similar boost inoculations performed on days 14 and 27. For primary or secondary immunizations with recombinant SARS-CoV-2 protein, a sterile 0.1 mL solution in PBS buffer containing 7.5-15 μg protein was mixed with an equivalent volume of emulsified Titermax Gold ® adjuvant immediately prior to administration. Viruslike particle display of mouse Fc-tagged RBD protein (Sino Biological) was accomplished using the T=4 VLP composed of 120 copies of a covalent dimer version of the PP7 coat protein with the ZZ-domain fused to the C-terminus, produced as previously described.^5^ This particle was incubated with mFc-antigen in a stoichiometric ratio of 3 ZZ-domains to 1 Fc domain (40 antigen molecules per particle) for 2 h at 37°C just prior to immunization. Complexation of Fc-antigen to the VLP was confirmed by dynamic light scattering (DynaPro, Wyatt Technology) on representative samples showing the expected increase in nanoparticle hydrodynamic radius. Blood was collected by submandibular bleeds on days 0 (pre-immunization), 14, 21, and 30 (terminal). Body mass was measured over time as a metric of general health and did not vary relative to vehicle-treated control mice. Mice selected for high anti-ectodomain serum antibody levels were sacrificed by CO2 asphyxiation on day 30 followed by splenectomy for B cell extraction. B cells were perfused from harvested spleens by aspirating 20 mL of Iscove’s Modified Dulbecco’s Media (IMDM) through tissue using 10-mL syringes and 22-gauge needles.

### Hybridoma fusion

SP2-IL6 myelomas (ATCC) cells were cultured in IMDM supplemented with antibiotics and 10% Low IgG FBS (Gibco). B-cells isolated from selected immunized BALB/c mice were incubated with 5 mL lysis buffer on ice for 5 minutes. After washing with 0.01M PBS, B cells were resuspended into serum free base IMDM and counted using a Cellometer K2 Fluorescent Viability Cell Counter (Nexcelom). The SP2 cells were then mixed with B cells in 1:1 ratio and centrifuged at 400 x g for 7 min. The cell pellet was resuspended in 2 mL serum-free base IMDM followed by addition of 50 μL of Pronase solution (10 mg/mL) for 3 min at room temperature. In this reaction, the total cell number varied between 5 x 10^7^ and 2 x 10^8^. 0.5 mL low IgG FBS was added to quench the reaction. After washing twice with Cytofusion medium C (BTX) by centrifugation at 400 x g for 7 minutes, the cells were resuspended in 10 mL Cytofusion medium C and loaded into a 10 mL flatpack chamber (BTX). Electrofusion using an ECM 2001 Electro Cell Manipulator with 630B Safety Stand (BTX) was performed under automatic mode for 5 minutes at a field strength of 3000 V/cm. The cells were then mixed with 40 mL pre-warmed medium and incubated in a 37°C water bath for 30 minutes before transferring into a T75 flask and incubated at 37°C with 5% CO2 for 48 h.

### Development of SARS2 anti-Spike mAbs

Fused hybridoma cells were plated into semi-solid methylcellulose-based media containing hypoxanthine, aminopterin, thymidine, and CloneDetect reagent. Automated selection using a ClonePix II instrument (Molecular Devices) isolated 350 IgG-positive colonies. Undiluted hybridoma supernatant from confluent wells was screened against 0.5 μg/mL r-spike ecto and 1 μg/mL His-S1 by indirect ELISA. Positive reactivity was scored if the average OD was greater than 2.0, a value far greater than negative-control signals obtained under the same conditions. In each plate run, mouse sera from the terminal bleed was included as a positive control. IgG-producing clones were cultured in static T-75s in IMDM with antibiotics and 10% Low IgG FBS at 37°C with 5% CO2. mAbs were purified by Protein G Sepharose Fast Flow and eluted at >0.2 mg/mL in 100 mM glycine/150 mM NaCl (pH 2.8) and neutralized with 200 mM Tris (pH 8.0). Antibodies were isotyped using the Rapid ELISA Mouse mAbs Isotyping Kit (ThermoFisher Pierce) with anti-mouse heavy chain capture antibody (anti-IgG1, IgG2a, IgGb, IgG3, IgA and IgM) or antimouse light chain (kappa or lambda) and analyzed by SDS-PAGE and Superdex-200 size-exclusion chromatography. Aliquots of mAbs were stored at 4°C until use. Protein concentrations were determined by NanoDrop 2000c spectrophotometer at 280 nm.

### Antibody sequencing

Hybridoma clones expressing anti-SARS-CoV-2 monoclonal antibodies were propagated in IMDM supplemented with antibiotics and 10% Low IgG FBS in static culture at 37°C with 5% CO2. 3-5 x 10^6^ Cells from each clone were harvested by centrifugation at 2,000 x g for 10 min and washed once in 0.01 M PBS (pH 7.4). Total RNA was extracted by RNeasy Mini kit (Qiagen) according to the manufacturer’s instructions.^29^ RNA reverse transcription and first strand cDNA synthesis were carried out using chain-specific reverse primers^30^ with ProtoScript® II First Strand cDNA Synthesis Kit (NEB). The resulting cDNA was used as template for preparing the sequencing library using 5’ multiplex PCR as previously described.^31^ The PCR products corresponding to the library amplification were purified on BluePippin with size cutoff around 500 bp. The gel purified amplification products were then indexed, purified, quantified, and validated according to instructions provided by the manufacturer (Illumina). The Illumina Version 3 (2 x 300 bp) sequencing kit was used to sequence the library, and sequencing data were processed using the MiXCR tool.^32^

### ELISA

Anti-spike immune responses, antigen specificity of immunized mouse sera, and antigen recognition of hybridoma supernatants were analyzed by ELISA. His-S1 or ecto-spike proteins were plated on half-area high-binding 96-well polystyrene plates (Corning) at 1 μg/mL or 0.5 μg/mL, respectively, in PBS overnight at 4°C. Unbound protein was washed away with PBST (1x PBS with 0.5% Tween) followed by blocking with 1% casein (w/v) in PBS (VWR International, Radnor, PA) for 2 h at room temperature with mild shaking (55 rpm). Serum or supernatant samples were then incubated for 1 hr at room temperature with shaking, followed by washing with PBST. For serum titers, six dilutions of sera in blocking buffer were plated; for supernatant specificity and epitope mapping experiments, undiluted hybridoma supernatant was plated. Secondary reporter goat anti-mouse IgG HRP (Southern Biotech) was diluted (1:2500) in blocking buffer and incubated for 1 h at room temperature with shaking, followed by washing. Plates were developed by adding 1-step Ultra TMB (Fisher Scientific) for 30 s (serum) or 90 s (supernatant), followed by quenching with 2N H2SO4. Absorbance (450 nm) was measured by platereader (Varioskan Flash, Thermo Fisher Scientific). Titers were calculated by sigmoidal non-linear regression using GraphPad Prism v.8 analysis with log 10 serum dilution plotted against absorbance at 450 nm.

### Spike detection sensitivity and specificity by indirect ELISA

Recombinant spike S1 or ectodomain proteins were plated at 0.5 ug/mL in 100 mM sodium bicarbonate (pH 9.0) in 96-well high-binding microtiter plates overnight at 4°C. The wells were washed with PBST, treated with blocking buffer (2% BSA in PBST) for 1 h at room temperature, and washed with PBST. Hybridoma supernatant or mAb (1 μg/ml) in blocking buffer was incubated for 1 h at room temperature, washed three times with PBST, and incubated with HRP-Goat anti-mouse IgG (1:5000) in blocking buffer (1 h, room temperature). After washing 3x with PBST, TMB substrate (100 μL, A and B, Sera Care) was added, followed after 5 min by 100 μL of stop solution (Sera Care). Absorbance was monitored at 450 nm on a Spectramax 384 microplate reader (Molecular Devices).

### Label-Free binding and kinetic studies

Label-free analysis of binding by biolayer interferometry (BLI) was performed on an Octet Red96 (Forte Bio) at 30°C in kinetics buffer (PBS, 0.05% BSA, 0.02% Tween20). Biosensors were equilibrated for 10 min in kinetics buffer and microplates filled with 200 μL of sample in kinetic buffer and agitated at 1000 rpm. For kinetic analysis of mAbs, anti-mouse Fc (AMC) biosensors were loaded with IgG (1-2 μg/mL) giving a loading response of 0.5-0.8 ± 0.15 nm among 8 sensors. The loaded sensors were exposed to r-spike ecto protein (15 μg/mL) with subtracted buffer reference for 1000 s and buffer dissociated for 2000 s in two independent assays (averaged values with R^2^>0.98). Data were analyzed in Octet Data Analysis 9.0. To deduce a direct binding affinity via the kinetic rate constants (1/K_ads_ = k_dis_/k_on_, where 1/K_ads_ = apparent equilibrium dissociation constant, kon = association rate constant, and koff = dissociation rate constant) the buffer-subtracted Octet data were fit locally (relative binding) or globally (kinetic analysis) to a simple 1:1 Langmuir model. Experiments performed on a subset of samples by immobilizing r-spike ecto (15 μg/mL) onto anti-His tag (HIS2) sensors gave similar results but with poorer-quality fits and so are not reported. Typical loading levels were 0.8-1.0 ± 0.15 nm within a row of eight tips, with standard deviation within the instrument noise. Purified IgG (100 nM) was associated for 150 s followed by dissociation in buffer for 200 s in two independent assays (R^2^>0.92) using a non-specific IgG reference.

### Epitope binning by BLI

Epitope binning was performed in tandem format using an Octet Red 96 instrument. Recombinant spike ecto protein in kinetics buffer was immobilized onto streptavidin (SA) biosensors (by virtue of the protein’s dual Strep-Tag® II peptides, protein concentration 10 μg/mL). Each epitope binning experiment consisted of a baseline measurement in kinetics buffer for 60 s; loading of r-spike ecto for 500 s; a second baseline in kinetics buffer for 60 s; association of the “primary” mAb for 400 s; dissociation in kinetics buffer for 300 s; association of the competing (“secondary”) mAb for 300 s; and a second dissociation in kinetics buffer for 200 s with shaking at 1000 rpm at 30°C. Binding response of the secondary mAb was measured against the saturating mAb. All mAb solutions were prepared in kinetics buffer. All of the data were analyzed using ForteBio Octet Data Analysis software version 10.0.

### Mix-and-read (split luciferase) RBD/ACE2 interaction assay

The ability of monoclonal antibodies to inhibit RBD/ACE2 interaction was quantified in a purpose-built homogeneous biochemical assay. Human codon optimized ACE2 (residues 1-615, NM_001371415) was fused to a C-terminal extension composed of a 5-residue linker, 8xHis, 16-residue linker and LgBit of Promega NanoBit split luciferase system. Receptor-binding domains of SARS-CoV-1 strain Urbani (AY278741, spike residues 1-14 and 306-527) and SARS-CoV-2 isolate Wuhan-Hu-1 (MN908947, spike residues 1-14 and 319-541) were fused at their C-termini to identical linkers and tag sequence with SmBit in place of the LgBit. These constructs were cloned into the episomal vector EEV600A-1 (System Biosciences) into which a puromycin resistance cassette was added so that cell pools expressing the proteins could be rapidly selected, if desired. Human Expi293 cells (Thermo Scientific) were transfected with these constructs using FectoPro (PolyPlus Transfection) and the proteins purified from culture supernatants by immobilized metal affinity chromatography using HisTrap Excel nickel columns (Cytiva). Purified proteins were dialyzed in PBS and stored at −80° C. The RBD/ACE2 assays were conducted 96-well plates with low-protein binding surface and 0.02% BSA in PBS (w/v) as the diluent. In the first step, 3.2 ng of RBD-SmBit proteins in 20 μL were mixed with 20 μL mAb dilution so that the mAb was at 20X molar excess over the RBD-SmBit. After 1 h incubation at room temperature, ACE2-LgBit was added in volume of 20 μL so that it was in 10X molar excess over the RBD-SmBit. After another 1 h incubation at RT, 60 μL of Promega NanoGlo reagent was added and luminescence quantified 10 min later with no-antibody wells serving as the baseline. The data represents average results from repeated, independent experiments (n = 2-7).

### BLI Assay of RBD-ACE2 binding inhibition

The results of the mix-and-read RBD/ACE2 interaction assay were corroborated by BLI. Samples comprised of a 1.4:1 molar ratio of mAbs (10 μg/mL) to mouse Fc-RBD (2.5 μg/mL, Sino Biological, 51.5 kDa) were complexed in kinetics buffer for 30 min at room temperature. The experiment was carried out at 30°C with agitation at 1000 rpm on an Octet Red 96 or an Octet KQe (Forte Bio). His2 biosensors were hydrated for 10 min in kinetics buffer, then loaded for 300 s with 2.5 μg/mL ACE2-His (residues 1 – 740) (Sino Biological) or truncated ACE2 (described above for the mix-and-read assay) in kinetics buffer. Pre-complexed samples were then associated for 300 s followed by dissociation in kinetics buffer for 300 s. Binding responses of mouse Fc-RBD to ACE2-His were measured with and without mAb present and were analyzed with Octet Data Analysis v. 9.0.

### Immunohistochemistry

Formalin-fixed paraffin-embedded lung samples were obtained from a previously confirmed fatal COVID-19 case at Infectious Diseases Pathology Branch, CDC, Atlanta. All samples and associated medical and autopsy records were provided in the context of diagnostic consultation, a routine public health service provided by CDC. As such, institutional review was not required for the testing described in this article. Cytoplasmic staining patterns were analyzed and compared with a commercial rabbit polyclonal SARS nucleocapsid antibody (Novus Biological) at 1:100 dilution, previously validated and used as a primary SARS-COV-2 IHC diagnostic assay.^22^ Immunohistochemical assays were performed using an indirect immunoalkaline phosphatase detection with the Mach 4 AP Polymer system (Biocare Medical). Briefly, 4 μm tissue sections were deparaffinized in xylene and rehydrated through graded alcohol solutions. Heat-induced epitope retrieval was performed using a citrate-based buffer and Decloaking Chamber (Biocare Medical). Primary antibody was applied at 1:100 for 30 min. The antibody-polymer conjugate was visualized by applying Permanent Red Chromogen to tissue sections for 30 min (CellMarque, MilliporeSigma). Slides were counterstained in Mayer’s Hematoxylin (Polyscientific), blued in lithium carbonate (Polysciences, Inc.) and cover slipped with aqueous mounting medium (Polysciences, Inc.)

Of the 13 antibodies tested, immunohistochemical staining of SARS-CoV-2 case control material was detected for two (1-3D2 and 1-3D7). The staining quality of each antibody was reliable and generally similar to the commercial polyclonal SARS-CoV-2 nucleocapsid antibody used as a positive control, the monoclonals showing similar staining intensity but fewer positive cells.

### Microneutralization assays

All SARS-CoV-2 microneutralization assays (MNT) were performed following biosafety level-3 precautions.

#### Using SARS-CoV-2 wild type isolate

The WA strain of SARS-CoV-2^17^ was employed using a modified version of a previously established protocol.^33^ Vero cell suspension (ATCC CCL-81) was prepared at 2.2 – 2.5 × 10^5^ cells/mL in DMEM (Thermo Fisher, catalog 11965118) + 10% fetal bovine serum (FBS, defined, Hyclone catalog SH30070.03, heat-inactivated 56°C for 30 min) + 2X antibiotic-antimycotic (Thermo Fisher catalog 15240062) + 2X penicillin-streptomycin (Thermo Fisher catalog 15140122) immediately before use. Purified monoclonal antibodies (50 μg/mL in serum-free DMEM) were 2-fold serial diluted in serum-free DMEM in a 96-well flat bottom plate, from 50 μg/mL to a 0.024 μg/mL, in triplicate, to a final volume of 50 μL/well. SARS-CoV-2 was diluted to a final working dilution of 700 TCID_50_/mL in serum free DMEM, and 50 μL was added to each well, giving final antibody concentrations ranging from 25 to 0.012 μg/mL. After 30 min incubation at 37°C and 5% CO2, 100 μL of Vero cells in suspension were added to each well, for a final concentration of 2.2 – 2.5 × 10^4^ cells/well. After 5 days further incubation at 37°C and 5% CO2 in a level-3 biosafety cabinet, media was aspirated from the wells, and 150 μL crystal violet fixative (0.15% crystal violet, 2.5% ethanol, 11% formaldehyde, 50% PBS, 0.01M pH 7.4) was added to each well. Plates were incubated for 20 min at room temperature. The fixative was aspirated, plates were washed with 200 μL/well distilled water and scored. The endpoint concentration at which antibodies were determined to be neutralizing for SARS-CoV-2 infection was the lowest concentration of antibody at which 3 replicate wells were protected against virus infection.

#### Using SARS-CoV-2 reporter virus

Vero E6 cells (ATCC CRL-1586) were suspended to a density of 2.2 x 10^5^ cells/mL in DMEM (Thermo Fisher catalog 11965118) with 10% FBS (Hyclone catalog SH30406.02HI) and 100 μL of the cell suspension was seeded into each well on flat-bottom 96-well cell culture plates. The plates were incubated at 37°C with 5% CO2 for 16-24 h to obtain a monolayer of confluent cells. Purified monoclonal antibody stock at a concentration of approximately 1mg/mL was 1:2 serially diluted in infection media (DMEM with 2% FBS and 1X penicillin-streptomycin (Thermo Fisher catalog 15140122) starting at 1:20 and ending at a dilution of 1:20,480. Dilutions were done with a final volume of 60 μL/well in quadruplicate rows using round-bottom 96-well plates. Recombinant mNeonGreen SARS-CoV-2 reporter virus icSARS-CoV-2-mNG^18^ was diluted to 2,000 TCID_50_/mL in infection media and an equal volume (60 μL) of the diluted virus was added to each well on the antibody dilution plates. Following 60 min incubation at room temperature, 100 μL of the virus-antibody mixture from each well was transferred to corresponding wells on the cell plates, from which the original cell culture media were decanted right before the transfer. After 2 days of incubation at 37 °C with 5% CO2, plates were scanned with the EVOS FL Auto Cell Imaging System to identify cells emitting green fluorescence. The endpoint dilution at which antibodies were determined to be neutralizing was the highest dilution of antibody at which more than half of the wells were absent of green fluorescence.

### Extended Data

Additional details regarding immunization, hybridoma generation and selection, *in vitro* characterization, sequencing, and competitive epitope binding are provided in the accompanying file.

## Acknowledgements

We are grateful to Dr. Barney Graham (Vaccine Research Center, NIAID) and Prof. Jason McClellan (University of Texas Austin) for the plasmid for the engineered spike ectodomain protein used in this work,^15^ under a material transfer agreement with the NIH. We also thank Jeff Noble for assistance with terminal mouse bleeds.

## Funding

This work was funded by the Centers for Disease Control and Prevention, and by the endowment for the James A. Carlos Family Chair for Pediatric Technology at the Georgia Institute of Technology.

## Author contributions

A. Chapman designed and performed all immunization schedules and assessed primary immune responses. J. Lee, A. Chida, and X. Tang prepared hybridoma fusions and selected initial clones. A. Chapman, A. Chida, J. Lee, and K. Mercer performed ELISA analyses. R. Wharton, K. Mercer, and J.M. Goldstein performed BLI analyses. M. Kainulainen, E. Bergeron, R. Wharton, and X. Tang produced the spike ectodomain protein. B. Zhou, A. Tamin, N. Thornburg and J. Harcourt performed neutralization assays. M. Kainulainen performed the split-luciferase ACE2 binding assay. B.C. Bollweb and R.B. Martines performed tissue immunohistochemical experiments. L. Zhao, A. Bryksin, X.Tang, and A. Chapman designed, performed, and analyzed mAb mRNA sequencing experiments. M. Schroeder, D. Wentworth, D. Petway and D. Bagarozzi, Jr. assisted in data management and analysis. A. Chapman, M. Finn, and J.M. Goldstein wrote the manuscript with input from all authors.

## Competing interests

None

## Disclaimer

The findings and conclusions in this report are those of the authors and do not necessarily represent the official position of the Centers for Disease Control and Prevention.

This activity was reviewed by CDC and was conducted consistent with applicable federal law and CDC policy.§ Footnote §See e.g., 45 C.F.R. part 46, 21 C.F.R. part 56; 42 U.S.C. §241(d); 5 U.S.C. §552a; 44 U.S.C. §3501 et seq.

## Notes

### Competing Interest Statement

The authors have declared no competing interest.

