## Extended Data for "Rapid Development of Neutralizing and Diagnostic SARS-COV-2 Mouse Monoclonal Antibodies"

‡ Authors provided equivalent contributions

* Corresponding authors

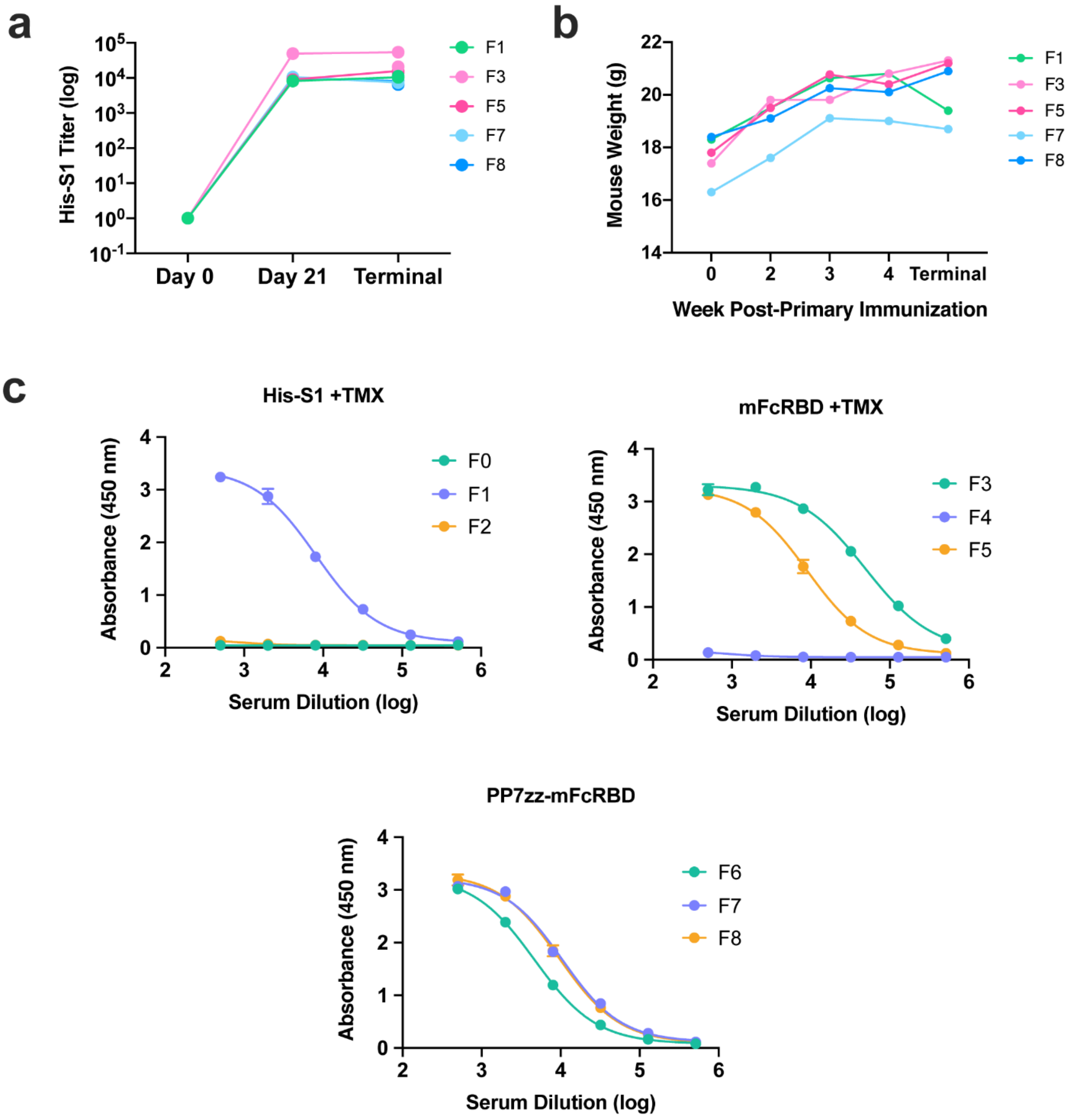


**Extended Data Figure 1. SARS-COV-2 Spike protein subunit vaccine safety and humoral immune response in mice. a**, Anti-S1 titers over time (fusion mice only) measured by ELISA against His-S1. Day 0 is preimmune sera, Day 21 is one week after the first boost, Terminal sera is Day 30. **b**, Fusion mouse weight over the course of vaccine schedule (n=5). **c**, Vaccine response at day 21 as assessed by ELISA. Immunized mouse sera from each group was collected, diluted, and tested against His-S1 (1 ug/mL) (F1: His-S1 + Titermax Gold® [Series 1]; F3 and F5: mFcS1-RBD + Titermax [Series 2]; F7-F8: PP7zzmFcS1-RBD + Titermax Gold ® adjuvant [Series 3]).

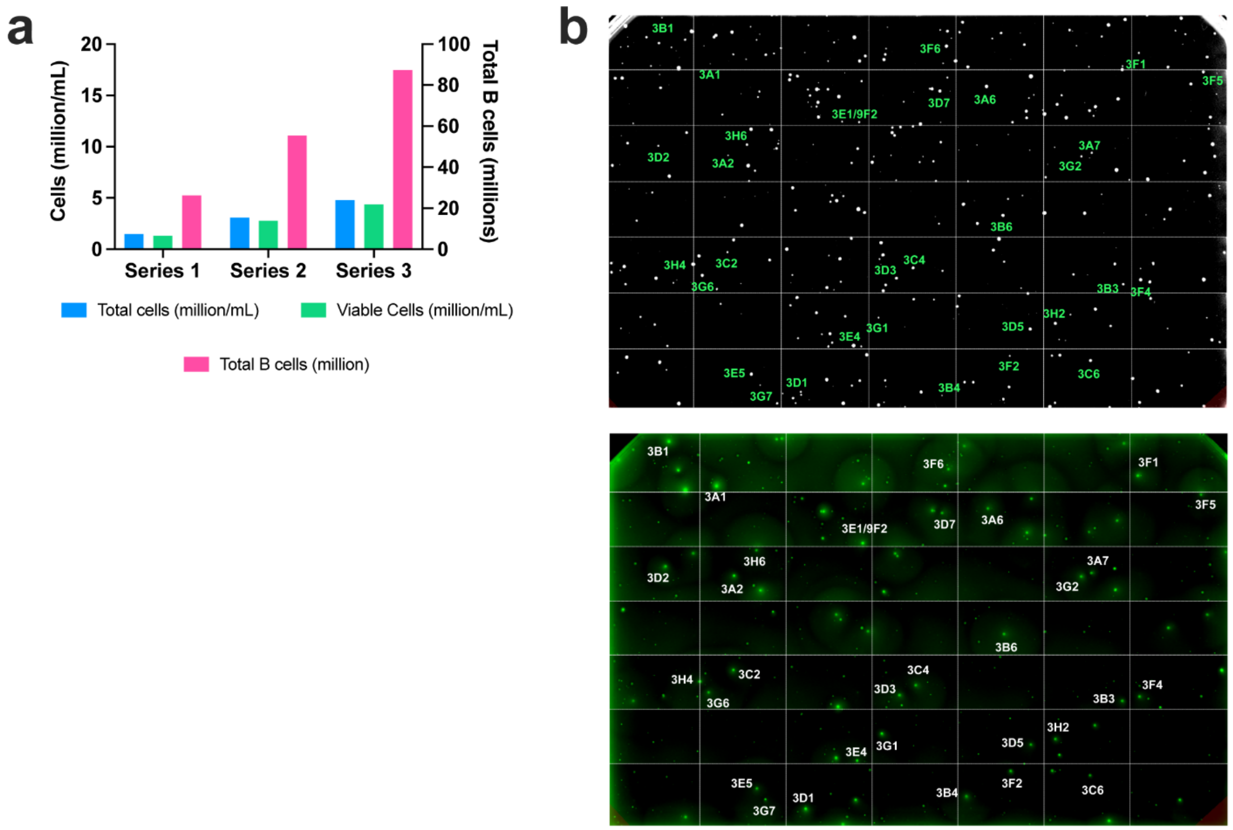


**Extended Data Figure 2. Hybridoma statistics from SARS-COV-2 immunized mice. a**, Total cell yield and B cell yield from mice immunized with either His-tag S1 (Series 1), mouse Fc-tagged S1 Receptor Binding Domain (Series 2), or codelivery of mFcRBD by complexation with PP7zz VLPs (Series 3), all in emulsion with Titermax Gold ®. **b**, Representative images from 3D culture of IgG secreting hybridoma from the combined B cells harvested from two mice immunized with mFcRBD. (*top*) brightfield, (*bottom*) anti-IgG FITC. Clones were chosen by ClonePix2 based on morphology of each clone and the diameter and brightness of the FITC halo, indicating the level of IgG secretion.

**Extended Data Table S1. Hybridoma statistics from SARS-COV-2 immunized mice.**

| [Fusion #], initial immunogen, (mice used) | Total B cells fused | Clones detected (fusion efficiency) | High/Low FITC clones detected (%) | Clones spike reactive (total clones picked) | Clones dual antigen reactive | Clones with ecto preference | Clones with S1 reactive preference |
|---|---|---|---|---|---|---|---|
| [1] SARS-COV-2 His S1 (mouse F1) | 25 x $10^6$ | 699 (2.8 x $10^{-5}$) | 16/304 (5.2%) | 27 (282) | 15 | 3 | 9 |
| [2] SARS-COV-2 mFcRBD (mice F3 + F5) | 55 | 667 (1.2 x $10^{-5}$) | 85/184 (46.1%) | 157 (258) | 107 | 48 | 2 |
| [3] SARS-COV-2 PP7zzmFcRBD + TiterMax (mice F7 + F8) | 75 | 307 (0.41 x $10^{-5}$) | 4/8 (50%) | 2 (12) | 1 | 1 | 0 |
| **Totals** | **155** | **1673 clones** | **105/496 (21.2%)** | **186 (552)** | **123** | **52** | **11** |

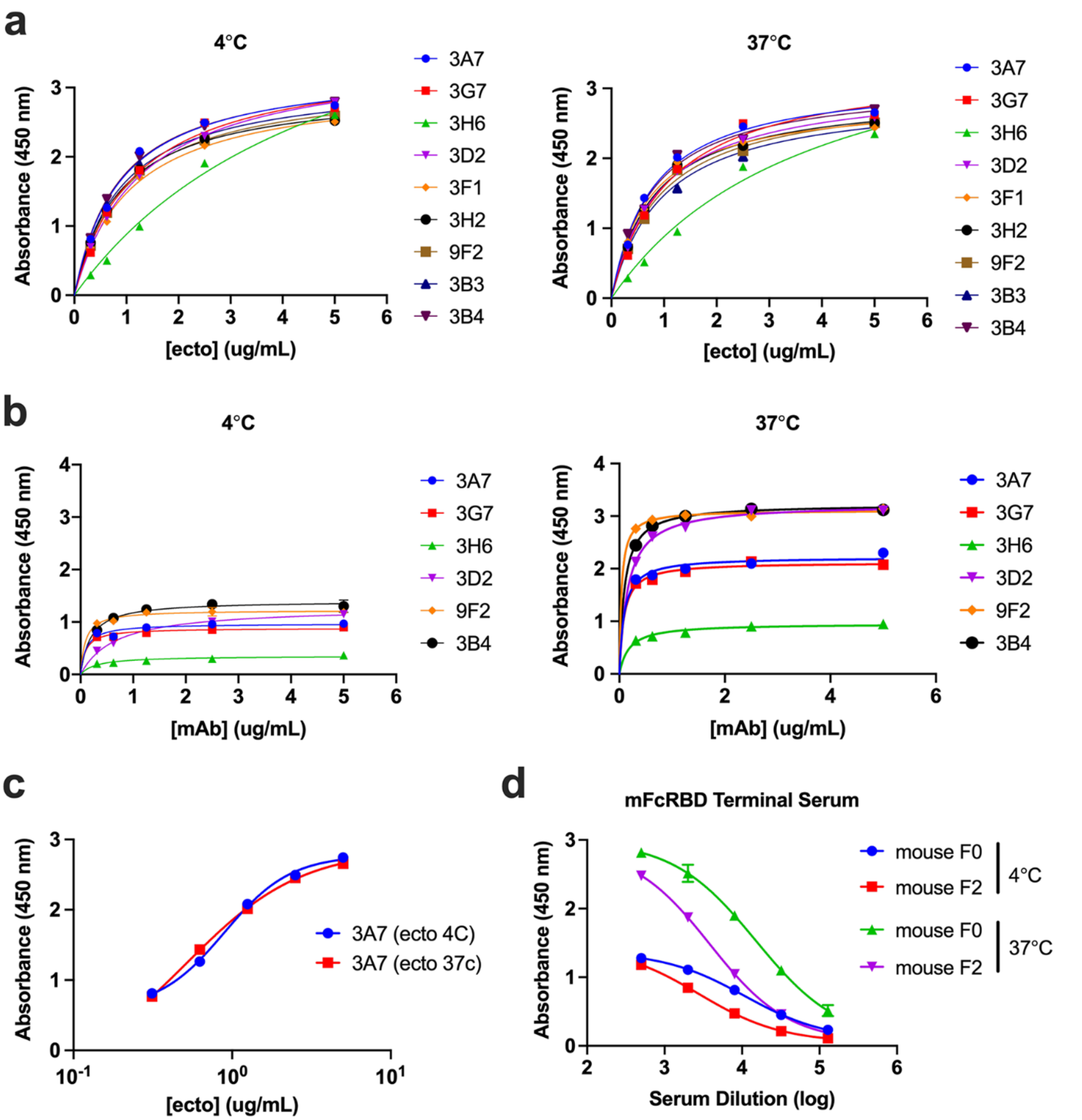


**Extended Data Figure 3. Thermal stability of SARS-CoV-2 spike ectodomain. a**, Decreasing concentrations of r-spike ecto protein were plated on streptavidin coated 96-well plates (5 ug/mL), with the protein either used directly from storage at 4°C or after 1 h incubation at 37°C. The indicated monoclonal antibodies were then introduced (10 μg/mL) and incubated for 1 h. After washing, goat anti-mouse IgG-HRP reporter (1:2000 dilution) was added for 1 h, and plates were developed with Ultra-TMB substrate (30 s) and quenched with 2N $H_2SO_4$. **b**, After equilibration at 4°C or 37°C for one hour, r-spike ecto was plated at 0.5 μg/mL for one hour followed by incubation with decreasing concentrations of select mAbs (5 μg/mL – 0 μg/mL) under the conditions described for panel (**a**). **c**, Comparative binding profile of anti-RBD mAb 3A7 to r-spike ecto plated after temperature equilibration at either 4°C or 37°C. **d**, Terminal sera ELISA from mice immunized with mFcRBD (from which 32 out of 33 selected mAbs originated) against temperature-varied r-spike ecto (0.5 μg/mL).

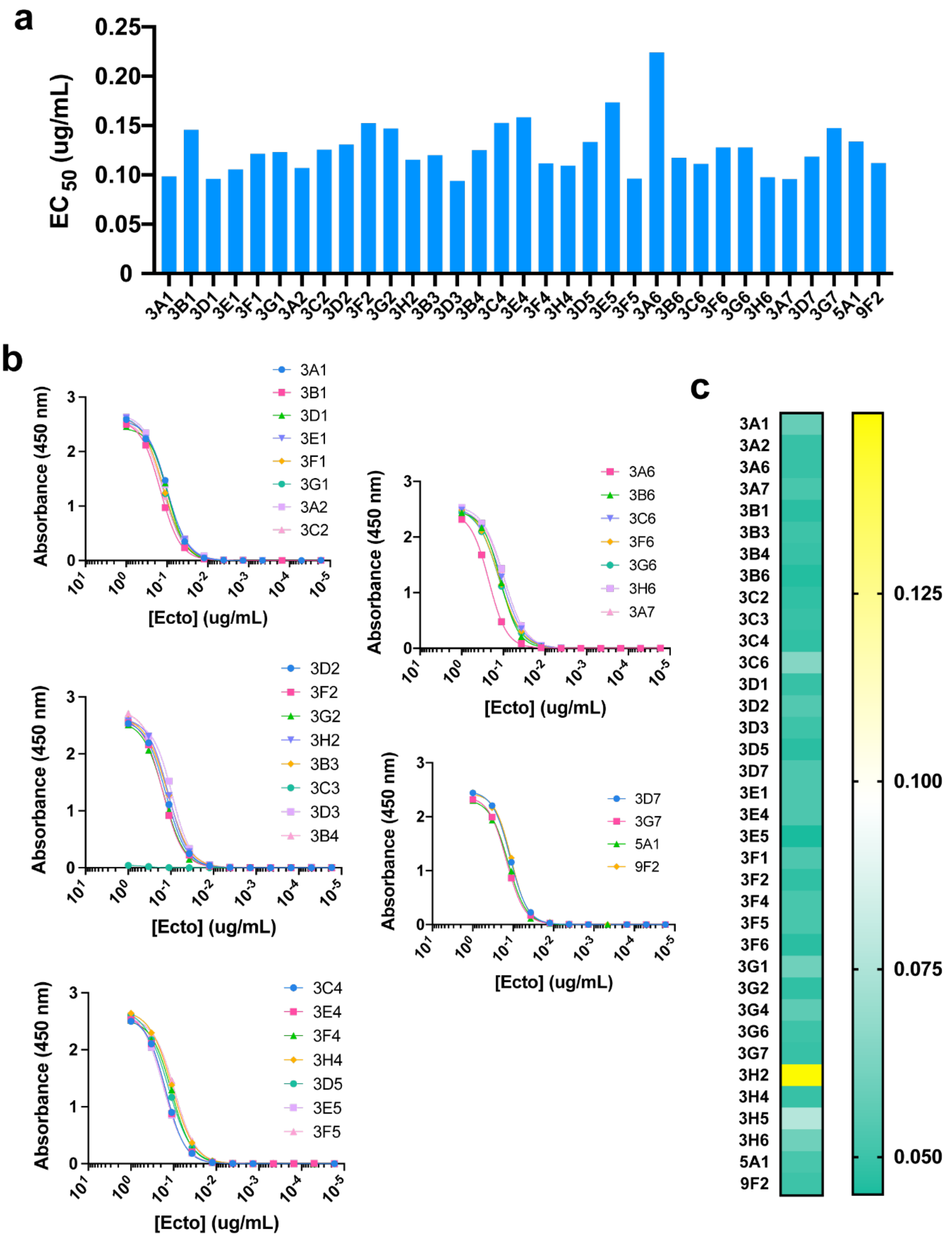


**Extended Data Figure 4. EC50 values for binding to plated r-spike ecto. a**, EC50 (μg/mL) of SARS-CoV-2 mAbs to trimeric spike ectodomain, calculated using EC50 shift nonlinear regression (GraphPad Prism v.8) from plots in panel (**b**). **b**, Indirect ELISA mAb (1 ug/mL) sensitivity to spike ectodomain (titration beginning at 1 ug/mL). **c**, Qualitative response (optical density) from indirect ELISA of mAbs against plated heat-inactivated SARS-CoV-2 (obtained from ATCC).

**Extended Data Table 2. Variable gene usage, CDR3 sequences and amino acid length of 32 sequenced SARS-CoV-2 monoclonal antibodies.** mAbs were sequenced by Illumina next generation sequencing, and heavy and light chain variable gene and CDR3 regions assigned by MiXIR software. Corresponding epitope bin is included, as determined by competitive BLI and summarized in Extended Figure 5.

| mAb | Epitope Bin | Sequence | | | | | |
|---|---|---|---|---|---|---|---|
| | | Heavy Chain | | | Light Chain | | |
| | | IGHV gene | CDR3 sequence | CDR3 length | IGKV gene | CDR3 sequence | CDR3 length |
| 3A2 | 1a | IGHV8-6 | CARRGPLITTDGTFDVW | 17 | IGKV6-23 | CQQYSSYPYTF | 11 |
| 3A7 | 1a | IGHV8-6 | CARRGPLITTDGTFDVW | 17 | IGKV6-23 | CQQYSSYPYTF | 11 |
| 3F5 | 1a | IGHV8-6 | CARRGPLITTDGTFDVW | 17 | IGKV6-23 | CQQYSSYPYTF | 11 |
| 3C6 | 1a | IGHV8-12 | CARRGPLITTDGTFDVW | 17 | IGKV6-13 | CQQYSSYPYTF | 11 |
| 3B3 | 1a | IGHV3-1 | CAGRRGAYYGNYEEDYW | 17 | IGKV4-55 | CQQWSSYPYTF | 11 |
| 3G7 | 1b | IGHV1-61 | CARSSGYDWYFDVW | 14 | IGKV3-5 | CQQSNEDPWTF | 11 |
| 3C2 | 1b | IGHV1-5 | CARSGGYDWYFDVW | 14 | IGKV3-5 | CQQSNEDPWTF | 11 |
| 3H6 | 2 | IGHV10-1 | CASYDGYRAWFAYW | 14 | IGKV1-117 | CSQSTHVPWTF | 11 |
| 3H2 | 3 | IGHV1-9 | CARNRFYWYFDVW | 13 | IGKV1-117 | CSQSTHVPWTF | 11 |
| 3F2 | 3 | IGHV1-12 | CARDGYFAMDYW | 12 | IGKV8-30 | CHQYYSYPWTF | 11 |
| 3E5 | 3 | IGHV1-20 | CGLRTYW | 7 | IGKV14-100 | CVQSVQFPYTF | 11 |
| 3G1 | 3 | IGHV2-2 | CAKYRYDSFAYW | 12 | IGKV3-12 | CQHSRELPYTF | 11 |
| 3E1 | 4 | IGHV14-1 | CARTYYYGSSYEAMDYW | 17 | IGKV6-32 | CQQDYSSPTF | 10 |
| 3D3 | 4 | IGHV14-1 | CARTYYYGSSYEAMDYW | 17 | IGKV6-32 | CQQDYSSPTF | 10 |
| 9F2 | 4 | IGHV14-1 | CARTYYYGSSYEAMDYW | 17 | IGKV6-32 | CQQDYSSPTF | 10 |
| 3H4 | 4 | IGHV14-1 | CARSYYDYDGGGCFDYW | 17 | IGKV6-32 | CQQGYSSPLTF | 11 |
| 3D7 | 4 | IGHV14-1 | CARSYYDYDGGGCFDYW | 17 | IGKV6-32 | CQQGYSSPLTF | 11 |
| 3B6 | 4 | IGHV14-1 | CARSYYTYDGFFDVW | 15 | IGKV6-32 | CQQDYSSPPTF | 11 |
| 3F6 | 4 | IGHV14-1 | CARSYYTYDGFFDVW | 15 | IGKV6-32 | CQQDYSSPPTF | 11 |
| 3F1 | 4 | IGHV14-1 | CTRYYYGSSGFFDVW | 15 | IGKV4-54 | CQQDYSSPTF | 10 |
| 3D5 | 4 | IGHV14-1 | CTRYYYGSSGFFDVW | 15 | IGKV3-5 | CQQDYSSPTF | 10 |
| 3B4 | 4 | IGHV14-1 | CARSYYYGTTSWFASW | 16 | IGKV13-85 | CQQYWSTPYTF | 11 |
| 3C4 | 4 | IGHV14-1 | CASGYDVNYELDYW | 14 | IGKV15-103 | CQQGQSYPYTF | 11 |
| 3E4 | 4 | IGHV14-1 | CASGYDVNYELDYW | 14 | IGKV15-103 | CQQGQSYPYTF | 11 |
| 3F4 | 4 | IGHV14-1 | CTRYYDYVYAMDYW | 14 | IGKV12-98 | CQQLYSTPLTF | 11 |
| 3D1 | 4 | IGHV14-1 | CVSGYYYYGSPYGAMDYW | 18 | IGKV14-111 | CQQYHGYPLTF | 11 |
| 3G2 | 4 | IGHV14-1 | CARWDFGNYVDYAMDYW | 17 | IGKV6-32 | CLQYDAFPWTF | 11 |
| 3G6 | 4 | IGHV14-1 | CARLNYDGYYDYAMDYW | 17 | IGHKV14-111 | CLQYDEFPFTF | 11 |
| 3D2 | 5 | IGHV1-4 | CARRYGNYDAWFTYW | 15 | IGKV6-23 | CQQYRTF | 7 |
| 3A1 | 6 | IGHV3-1 | CARWGNGKNAMDYW | 15 | IGHV12-44 | CQHHYGSPPTF | 11 |
| 3A6 | 6 | IGHV2-9 | CGRDYGILLIDYW | 12 | IGKV4-63 | CFQGSGYPFTF | 11 |
| 3B1 | 6 | IGHV3-6 | CARVDYDVGHWFAYW | 15 | IGKV4-50 | CHQFTTSPWTF | 11 |
| 5A1 | 6 | not sequenced | | | not sequenced | | |

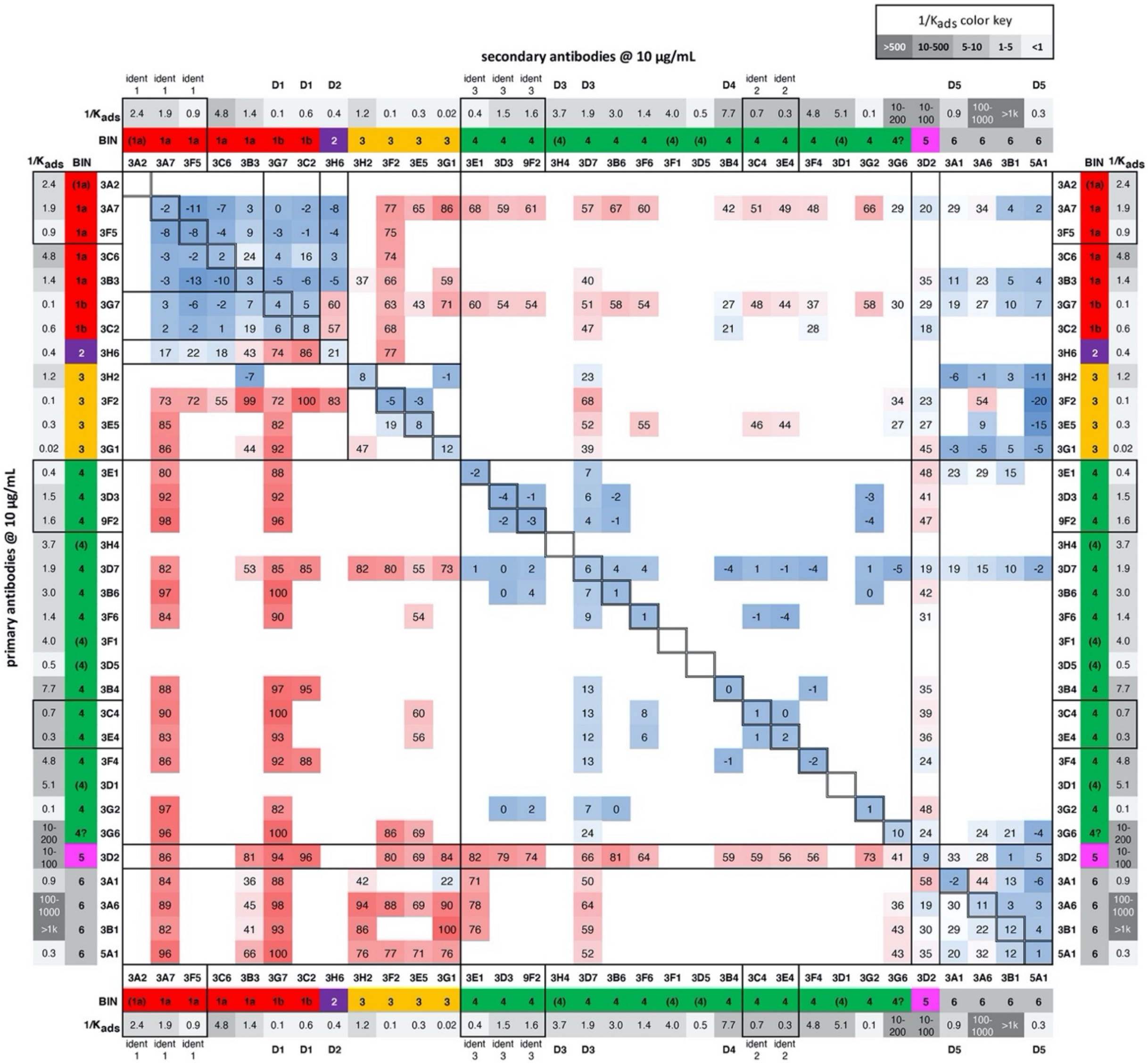


**notable disconnections between binding and split-luciferase results**

| class | mAbs | label | binding to full-length ACE2 | binding to truncated ACE2 | split luc SARS-1 | split luc SARS-2 |
|---|---|---|---|---|---|---|
| 1b | 3G7,3C2 | D1 | mod. Inhib | enhanced | little effect | little effect |
| 2 | 3H6 | D2 | inhib | inhib | little effect | little effect |
| 4 | 3H4,3D7 | D3 | little effect | enhanced | little effect | little effect |
| 4 | 3B4 | D4 | mod. Inhib | enhanced | enhanced | enhanced |
| 6 | 3A1,5A1 | D5 | little effect | little effect | little effect | mod-strong inhib |

**Extended Data Figure 5.** Collected from competitive binding experiments of the type shown in Fig. 3f and Extended Fig. 6, performed on groups of up to 13 mAbs at a time. BLI values for each experiment were normalized to a maximum of 100 to allow for incorporation into this master dataset. Color-coded bin assignments are the same as done by sequence. When rows do not match columns, the avidities of the antibodies were usually significantly different, skewing the results of this competitive binding assay. Blank squares denote comparisons not made. Most of group 6 (5A1, 3B1, 3G4, 3C3, 3H5, 9B1) are included because some data was collected, but they were not sequenced and are omitted from the list of 33 selected antibodies discussed in the main text. The three sets of identical antibodies are denoted by “ident #”. The table at the bottom summarizes results described in the text concerning the experiments shown in Figure 4a vs. 4f, highlighting the connection between competitive binding, functional properties, and sequence.

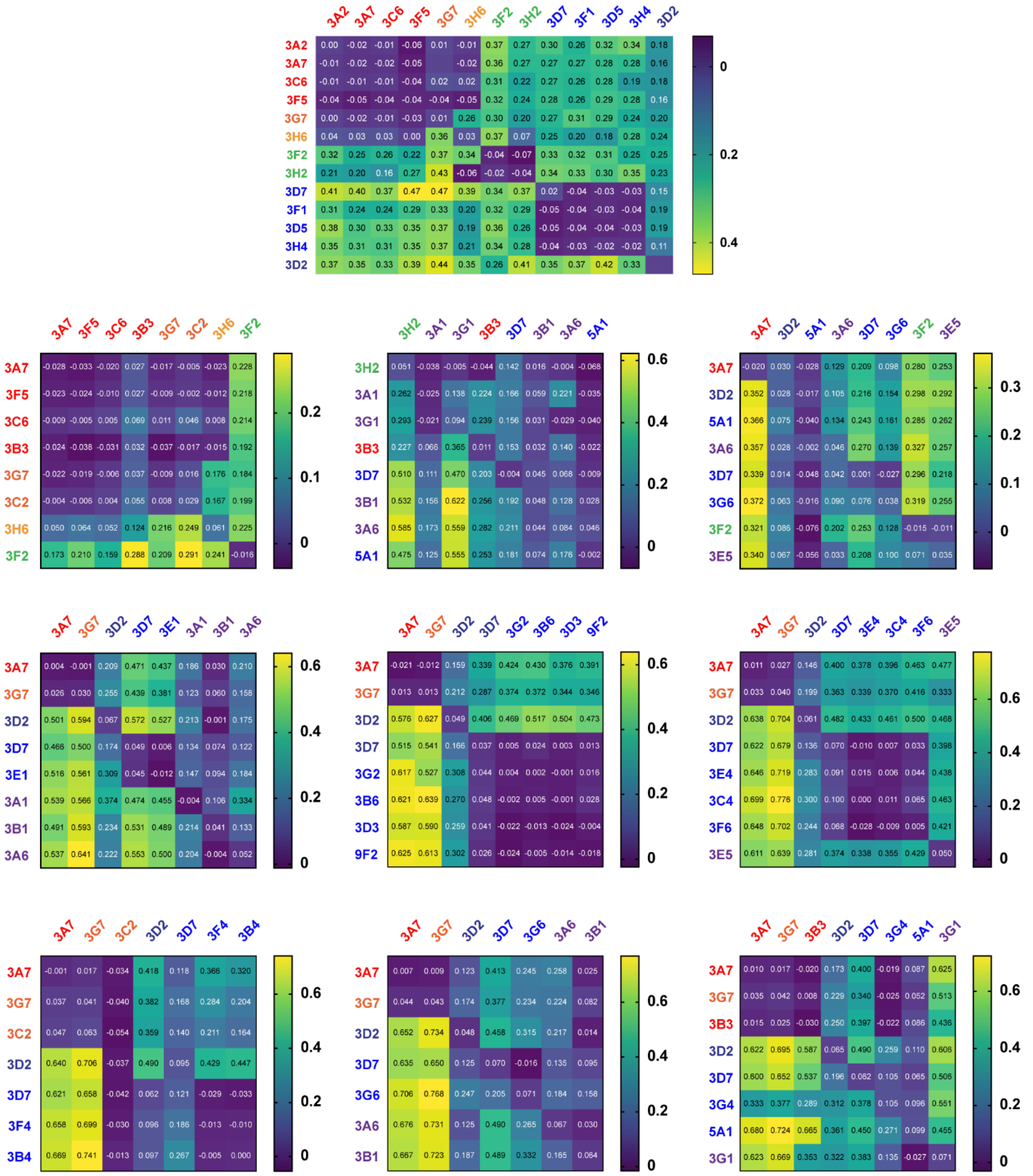


**Extended Data Figure 6. Individual epitope binning results**. Epitope classification experiments performed by BLI. r-Spike ectodomain protein (bearing twin Strep-Tag II sequences) was immobilized on streptavidin-coated biosensors (10 μg/mL). Primary antibody (y axis) was incubated for 400 s, allowed to dissociate in buffer for 300 s, followed by incubation with secondary antibody (x axis) for 300 s. Data analyzed using ForteBio Octet Data Analysis software v. 10.0.
